# Metagenomic estimation of absolute bacterial biomass in the mammalian gut through host-derived read normalization

**DOI:** 10.1101/2025.01.07.631807

**Authors:** Gechlang Tang, Alex V. Carr, Crystal Perez, Katherine Ramos Sarmiento, Lisa Levy, Johanna W. Lampe, Christian Diener, Sean M. Gibbons

## Abstract

Absolute bacterial biomass estimation in the human gut is crucial for understanding microbiome dynamics and host-microbe interactions. Current methods for quantifying bacterial biomass in stool, such as flow cytometry, qPCR, or spike-ins (i.e., adding cells or DNA from an organism not normally found in a sample), can be labor-intensive, costly, and confounded by factors like water content, DNA extraction efficiency, PCR inhibitors, and other technical challenges that add bias and noise. We propose a simple, cost-effective approach that circumvents some of these technical challenges: directly estimating bacterial biomass from metagenomes using bacterial-to-host (B:H) read ratios. We compare B:H ratios to the standard methods outlined above, demonstrating that B:H ratios are useful proxies for bacterial biomass in stool and possibly in other host-associated substrates. We show how B:H ratios can be used to track antibiotic treatment response and recovery in both mice and humans, which showed 403-fold and 45-fold reductions in bacterial biomass during antibiotic treatment, respectively. Our results indicate that host and bacterial metagenomic DNA fractions in human stool fluctuate longitudinally around a stable mean in healthy individuals, and the average host read fraction varies across healthy individuals by < 8-9 fold. B:H ratios offer a convenient alternative to other absolute biomass quantification methods, without the need for additional measurements, experimental design considerations, or machine learning algorithms, enabling retrospective absolute biomass estimates from existing stool metagenomic data.

## Introduction

The mammalian gut is a diverse and dynamic ecosystem, comprising microorganisms from all domains of life, including archaea, bacteria, viruses, and eukaryotes ^1^. Bacteria are the most abundant microbes in the gut, by mass, reaching densities of 10^11^ – 10^12^ cells per gram of stool and making up between 25-54% of stool dry weight ^2,3^. The gut microbiota confers essential biomolecular functions to the host ^4,5^. Disruption to this ecosystem, as in the case of antibiotic treatment ^6^, can increase susceptibility to opportunistic infections and other diseases ^2,4^. We know that the composition of the gut microbiota is shaped by a combination of intrinsic and extrinsic host factors, such as host genotype, physiology, immunity, behavior, and diet ^7–9^. Diet and behavior appear to exert the strongest influence ^8^. Gut microbiome composition is commonly quantified using shotgun metagenomic sequencing of fecal DNA, which provides relative, but not absolute, abundance estimates for microbial taxa and genes ^10^. Prior work has suggested that accurate estimates of absolute abundances in the gut are crucial to fully understanding cross-sectional and longitudinal variation in this important ecosystem ^11–14^.

Metagenomic shotgun sequencing is a cost-effective approach to comprehensively quantifying the ecological composition and functional potential of the gut microbiome ^2,15^. However, standard methods for quantifying absolute biomass require additional measurements beyond the metagenome ^16,17^. For example, flow cytometry of dilute stool homogenates can be used to estimate the number of cells per gram of feces ^18^. Cytometry can be labor intensive, requiring a dedicated cytometer and extensive standardization, in part due to the large amount of non-cellular debris and caustic compounds present in stool. Additionally, qPCR can be leveraged to detect the total copy number of the 16S gene per gram of stool (or some other marker gene), but PCR can be noisy and sensitive to inhibitors that are common in stool homogenates ^19^. Furthermore, qPCR absolute abundances are related to fecal DNA extraction efficiency (e.g., samples with lower extraction efficiency will appear to have lower biomass, independent of microbial load) ^17,20^. Spike-ins of DNA (post-extraction) or cells (pre-extraction) from organisms that are not normally present in the system can be used to renormalize relative gut bacterial abundances and obtain absolute biomass estimates ^16^. Spike-ins are excellent solutions to absolute biomass quantification, but they require additional sample processing steps and result in a reduced number of reads derived from the sample. Finally, a recent approach leverages machine learning to predict microbial load in stool metagenomes directly from bacterial taxonomic profiles, but this method relies on cytometric biomass estimates as the gold-standard and achieved somewhat marginal correlation coefficients with out-of-sample microbial load estimates (R=0.5-0.6) ^21^. All of the biomass estimates listed above are calculated per unit wet-weight (i.e., weight of a fresh sample, including the weight of the water in that sample), as opposed to dry-weight (i.e., weight is taken before and after drying the sample in an oven, so water weight can be subtracted), so that these microbial biomass estimates are generally conflated with fecal water content ^17,20^. However, total fecal biomass in the gut may vary independently of fecal water content. In summary, standard methods for estimating absolute biomass require additional measurements, experimental design considerations, and can suffer from confounding and bias.

Alternatively, some have argued for the use of standard reference frames applied directly to compositional data ^10^. Specifically, methods have emerged that use log-ratios of different microbiome features to break the underlying compositionality of the data and circumvent the need to estimate total microbial biomass ^10,22^. These methods are not dissimilar from spike-ins (e.g., dividing one value by another and taking the log), but leverage features that are already measured in the context of the metagenomic data. The downside to many of these log-ratio methods is that the resulting features become more difficult to interpret. In the simplest case of an additive log ratio, a single taxon is used to normalize the relative abundances of other taxa in the sample, but it is unclear what common denominator taxon should be used as a ‘control’. Here, we propose using the relative abundance of host DNA as that common denominator in stool metagenomic data, dividing the number of bacterial read counts by the number of host read counts to generate a bacteria-to-host (B:H) ratio. The key assumption is that the average rate of host DNA shedding into stool is relatively constant within and across healthy individuals, allowing us to treat host DNA as a naturally occurring, systemic spike-in. However, the degree to which this assumption holds true across a range of conditions will require scrutiny. Here, we compare ln(B:H) ratios to paired biomass measures derived from flow cytometry, qPCR, and synthetic spike-ins from a number of published studies ^4,16,23,24,25^. We assess whether or not B:H ratios are associated with fecal water content or stool consistency. We look at the variation in B:H ratios within and across healthy individuals. Finally, we assess how well stool B:H ratios capture known bacterial biomass trajectories following antibiotic treatment in both mice and humans. We conclude that normalization by host read fraction provides a useful estimate of absolute bacterial biomass in stool metagenomes from healthy individuals, without the additional expense or effort of flow cytometry, qPCR, or synthetic spike-ins.

## Results

### Confounding between bacterial load and stool water content

Estimates of gut bacterial biomass are often made per unit wet weight (e.g., cells or 16S copies per gram of fresh stool), unlike bacterial biomass estimates in many other systems, such as soils, which are often normalized to grams dry weight ^26,27^. Fresh stool samples can vary substantially in water content, depending on intestinal transit time, with constipation associated with reduced water content and diarrhea associated with elevated water content ^28^. The Bristol stool scale provides an ordinal score representing stool consistency, with lower scores (1-2) representing hard stools and higher scores (6-7) representing loose stools ^29^. Bacterial cell density per unit wet weight has been observed to be inversely proportional to stool water content ^30^, which suggests that current standard estimates of gut bacterial biomass are conflated with water content and intestinal transit time^21^. Let us imagine a scenario where the total bacterial biomass of a given bowel movement remains fixed, while water content and total stool mass can vary along the Bristol scale (**Fig. 1A**). If a standard amount of fresh stool (e.g., 200 mg) is sampled for biomass quantification (e.g., cytometry or DNA extraction), there will be an inverse relationship between cell count or 16S copy number and water content (**Fig. 1B-C**). Synthetic spike-ins are often applied at the aliquot level (e.g., to 200 mg of stool), which introduces this same water content confounding (**Fig. 1D**). In the mammalian gut, on the other hand, we postulate that epithelial cells are continually shed into stool as it passes throughout the system, resulting in a naturally-occurring spike-in, which may be independent of fecal water content, that could potentially be used to approximate the total bacterial biomass within the entire length of the colon, (**Fig. 1D**).

**Figure 1.**
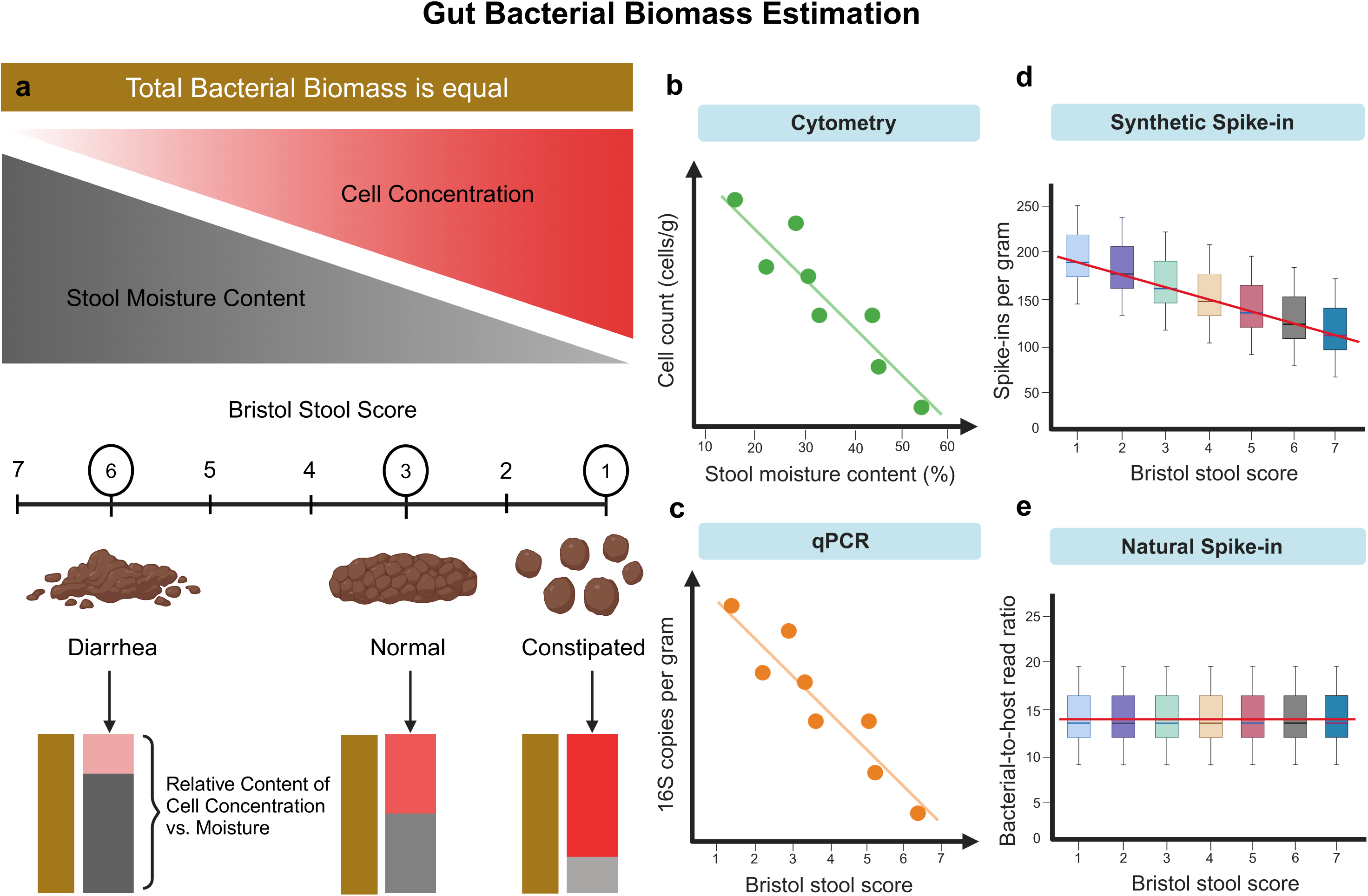
Conceptual figure explaining the potential confounding between stool moisture content and biomass estimates. (**A**) A schematic of how moisture content and cell count per gram are related, assuming constant biomass. Here, stool consistency is measured by the Bristol stool score (i.e., a proxy for water content), ranging from 1 (constipated; low moisture) to 7 (diarrhea; high moisture). Constipated stools (1-2) exhibit a high density of bacterial cells per gram but contain low moisture, while loose stools (6-7) have higher moisture levels with lower bacterial cell density per gram. (**B**) In this cartoon example, stool moisture content is inversely associated with cell concentration (cells/g of fresh or frozen stool), even when total bacterial biomass per gram dry mass is constant. (**C**) Biomass estimates derived from qPCR measurements from DNA extractions performed on an aliquot of fresh or frozen stool are also confounded with water content (**D**) Spike-ins can also be confounded by water content, if the spike-in is added to a specified aliquot of fresh or frozen stool. (**E)** However, if the spike-in is mixed into the entire bolus of stool within a person’s gut, like in the case of host DNA (i.e., a ‘natural spike-in’), this moisture confounding is no longer an issue.

**Table 1.** Summary of datasets used in this study.

| Dataset | Number of Subjects | Number of Samples | Sample Type | Data Type |
| --- | --- | --- | --- | --- |
| *Vandeputte et al.*<br>(2017) | 40 | 40 | Fecal samples | 16S amplicon data,<br>Cytometry; stool<br>moisture content |
| *Fromentin et al.*<br>(2022) | 1241 | 1680 | Fecal samples | Metagenomic data;<br>cytometry |
| Gut Puzzle Study<br>(this paper)<sup>1</sup> | 39 | 39 | Fecal samples | Metagenomic data;<br>Bristol stool score |
| *Chng et al.*<br>(2020) | 27 | 243 | Fecal samples | Metagenomic data;<br>qPCR |
| *Wallace et al.*<br>(bioRxiv, 2023)<sup>1</sup> | N/A | 385 | Cow milk samples | Metagenomic data;<br>Spike-in |
| *Palleja et al.*<br>(2018) | 12 | 58 | Fecal samples | Metagenomic data |
| *Poyet et al.*<br>(2019) | 4 | 398 | Fecal samples | Metagenomic data |

**Footnote:** <sup>1</sup> Number of subjects not reported for this dataset

### Bacteria-to-host (B:H) ratios are weakly associated with cytometric biomass measures, but not with stool consistency measures

We pulled down existing data from Vanderputte et al. (2017), which included paired measures of cytometric bacterial biomass estimates and moisture content from 223 stool samples ^18^. As expected, we observed a significant inverse association between cytometric bacterial biomass estimates (cells per gram of stool) and percent stool moisture content (linear regression, r^2^ = 0.14, P < 0.001; **Fig. 2A**). We were not able to calculate B:H ratios for the same set of samples because Vanderputte et al. generated 16S amplicon sequencing data, rather than metagenomes. However, in a larger metagenomic dataset of 1,883 stool samples from the MetaCardis cohort, excluding 203 samples with missing cytometric biomass values, an extremely subtle, but significant, positive association was observed between ln(B:H) ratios and cytometric bacterial biomass estimates (linear regression, r^2^ = 0.005, P = 0.005; **Fig. 2B**). MetaCardis did not include moisture content measures, so in order to obtain paired measures of stool consistency (Bristol stool scores; a proxy for stool water content) and B:H ratios, we generated new data from 39 healthy stool donors (**Fig. 2C**). We saw no significant association between the log B:H ratios and Bristol scores (ordinal logistic regression, P = 0.441), nor between the human read fraction and Bristol scores (ordinal logistic regression, P = 0.418; **Fig. S1)**. Overall, we find that cytometric measures of bacterial biomass show a negative association with moisture content and a weak positive association with B:H ratios (**Fig. 2A-B**). We did not find evidence for the same kind of association between B:H ratios and stool consistency, which is consistent with our hypothesis that B:H ratios are more direct estimates of absolute gut bacterial biomass, and less conflated with fecal water content.

**Figure 2.**
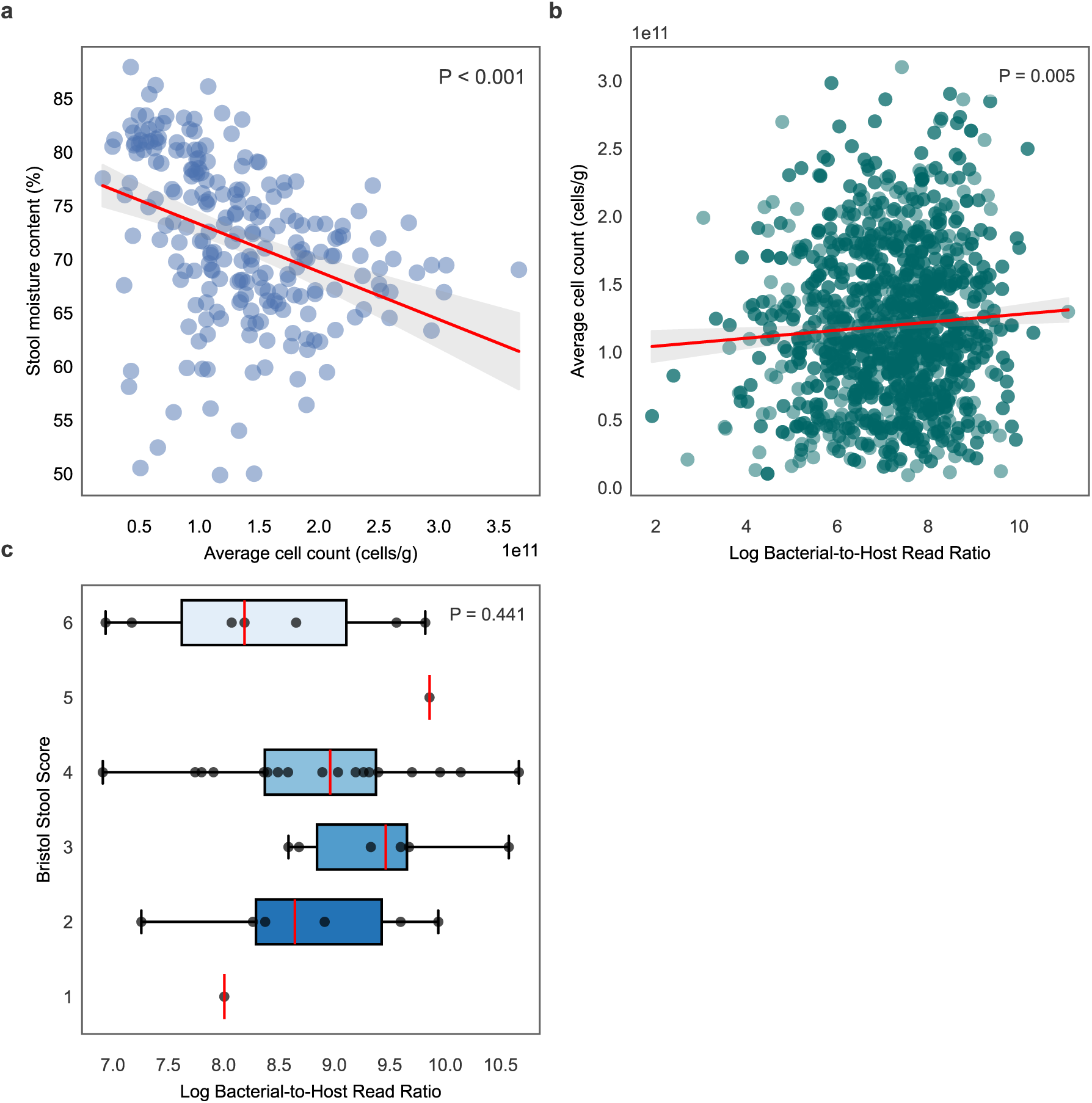
Comparing cell counts per gram (microbial load), moisture content, and B:H ratios. (**A**) Scatterplot shows a statistically significant negative association between bacterial cell counts per gram of wet stool and stool moisture content (n = 223). The red line represents the linear regression fit (R^2^ = 0.140, P < 0.001) of data obtained from Vandeputte et al. (2017). (**B**) Scatterplot shows a weak, but statistically significant, positive association between log-transformed B:H ratios and bacterial cell counts per gram of wet stool (n = 1680). The red line represents the linear regression fit (R^2^ = 0.005, P = 0.005) of data derived from Fromentin et al. (2022). (**C**) Boxplots showing B:H ratios across Bristol stool score categories (n = 39). Each boxplot displays the center line (median), box limits (first and third quartiles), and whiskers (1.5 × interquartile range). Using ordinal logistic regression, we did not observe a significant association between B:H ratios and Bristol scores (P = 0.441).

### B:H ratios in mice show quantitative agreement with qPCR and dietary-read-based biomass normalization

In data obtained from Chng et al. (2020) ^4^, we found that log-transformed B:H ratios in mice, derived from shotgun metagenomic data, were significantly associated with log-transformed absolute 16S rRNA genes copies quantified by qPCR across 107 fecal pellets (r^2^ = 0.656, P < 0.001; **Fig. 3A**). We also observed a significant association between log-transformed B:H ratios and total bacterial biomass estimates normalized by log-transformed bacterial-to-diet read ratios from metagenomic sequencing data, across 242 mouse fecal pellets from the same study (r² = 0.718, P < 0.001; **Fig. 3B**). In summary, we see strong agreement between B:H ratios, qPCR-based bacterial biomass estimates, and bacteria-to-dietary read ratios (i.e., normalized to plant-derived reads, which are likely from the diet; this diet normalization was reported as an alternative absolute biomass estimation approach in the Chng et al. paper) in mouse stool.

**Figure 3.**
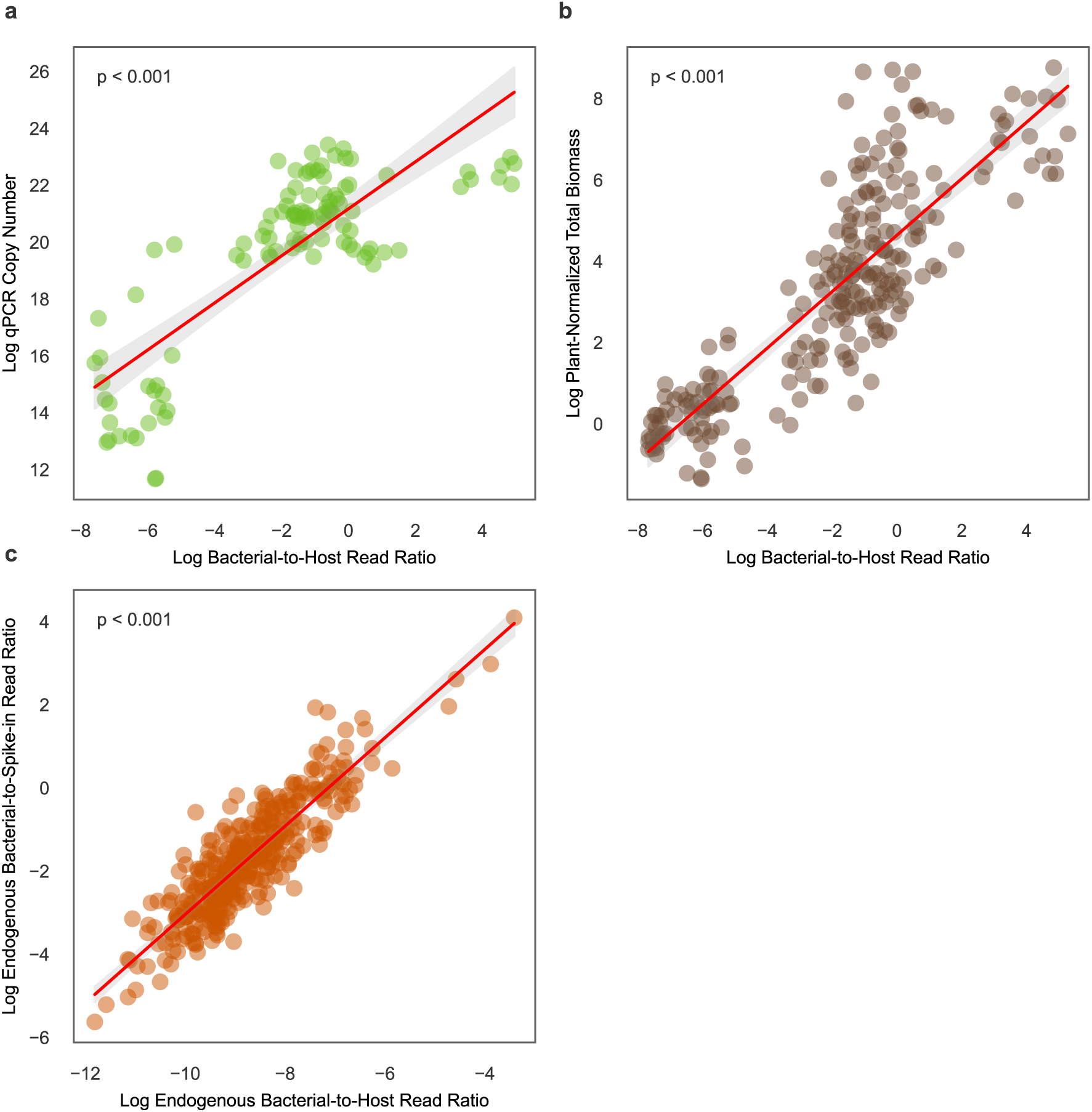
Associations between B:H read ratios, 16S qPCR-based bacterial biomass estimates, diet-read normalized bacterial biomass estimates, and spike-in normalized bacterial biomass estimates. (**A**) Scatterplot depicting a significant positive association between log-transformed B:H read ratios and qPCR-quantified biomass in mouse stools (log 16S copy number per microliter; n = 107). The red line represents the linear regression fit (R^2^ = 0.656, P < 0.001). (**B**) Scatterplot depicting a significant positive association between log-transformed B:H read ratios and shotgun sequencing-based diet-normalized total bacterial biomass estimates in mouse fecal samples (normalized by plant-derived reads present in the stool metagenome; n=242). The red line represents the linear regression fit (R^2^ = 0.718, P < 0.001). (**C**) Scatterplot illustrates a significant positive association between log-transformed B:H read ratios and log-transformed endogenous-to-spike-in ratios total bacterial reads from metagenomic sequencing of cow milk samples (n = 385). The red line represents the linear regression fit (R^2^ = 0.784, P < 0.001). Data for (**A**) and (**B**) were obtained from Chng et al. (2020), and data for (**C**) were obtained from Wallace et al. (2023).

### Comparing synthetic and natural spike-ins for biomass estimation in milk metagenomes

We had trouble identifying an appropriate stool spike-in dataset. As an alternative, we pulled down data from Wallace et al. (2023), encompassing metagenomic data from 385 cow milk samples with controlled bacterial spike-ins (i.e., a specific microbe that was known to be absent from milk). We observed a strong positive association between log-transformed B:H ratios and log-transformed total endogenous bacteria-to-spike-in ratios (r^2^ = 0.784, P < 0.001; **Fig. 3C**). This robust association supports the concept that host reads serve as a naturally-occurring spike-in that can be leveraged to estimate absolute bacterial biomass in metagenomic data sets from other host-associated substrates beyond stool.

### Examining intra- and inter-individual variation in bacterial and human relative DNA abundances in stool from healthy individuals

In data obtained from Poyet et al. (2019) ^31^, we were able to observe day-to-day fluctuations in B:H ratios, bacterial relative abundances, and host-read relative abundances across four individuals with long, dense stool metagenomic time series (**Fig. 4A-F**). Inter-individual differences in the mean ln(B:H) were significant between all donors, except for between donors am and ao (two-side Welch’s T-test, p < 0.001; **Fig. 4B**). Similar patterns were seen for the relative abundances of human and bacterial reads across these four donors (**Fig. 4C-F**). The largest fold difference in average B:H ratios across these four healthy donors was 2.4, between donors an and am (**Fig. 4B**).

**Figure 4.**
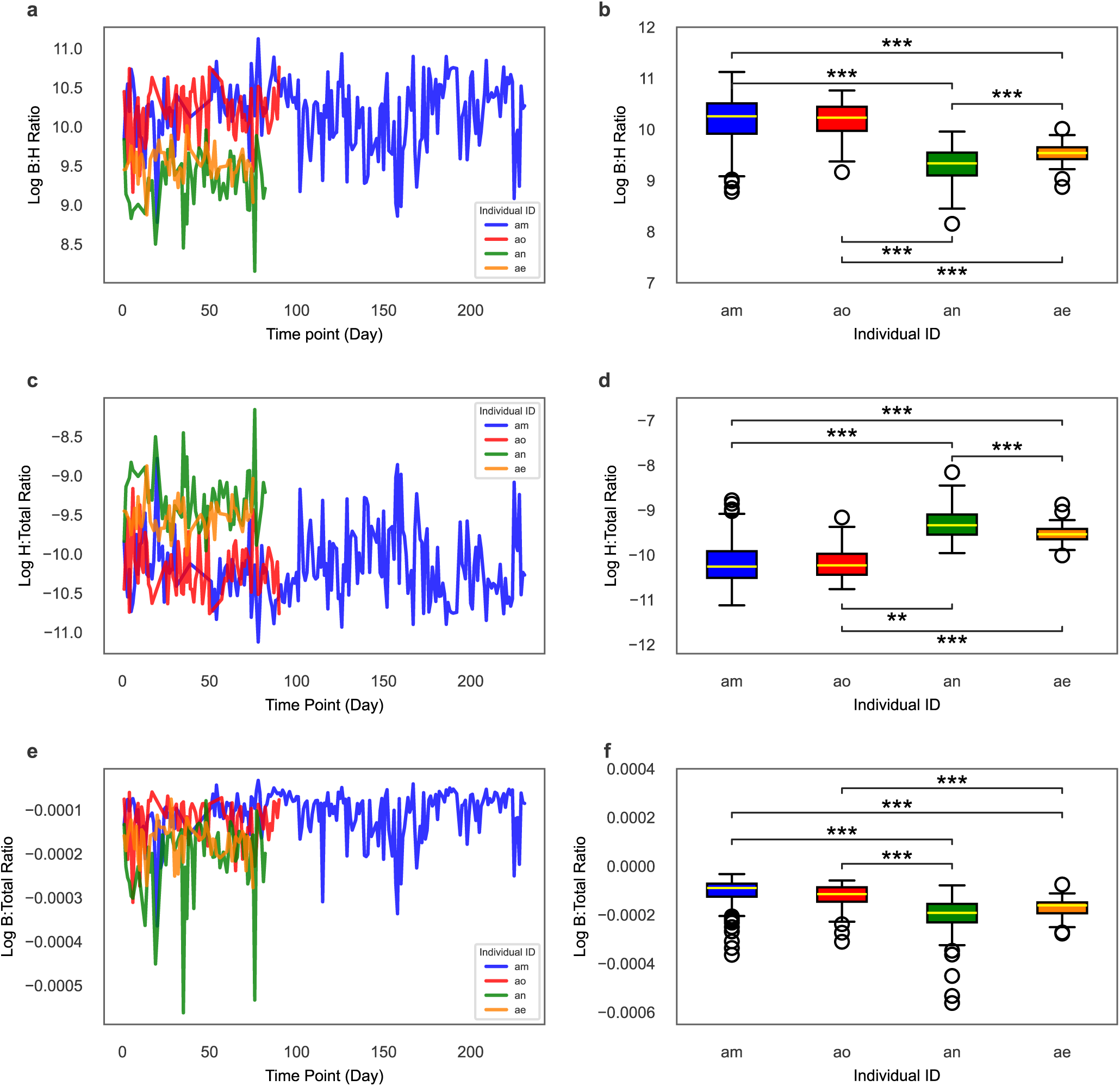
Temporal dynamics of bacteria-to-host ratios in healthy humans. (**A**) Dense time-series plot displaying an overall trend of stable, natural day-to-day intra-individual fluctuations in log-transformed B:H read ratios, derived from metagenomic sequencing of human stool samples collected from four healthy individuals (n=205 for donor am; n=57 for donor ae; n=62 for donor an; and n=74 for donor ao). (**B**) Boxplot showing the distributions of log-transformed B:H read ratios across the same four healthy individuals. Each boxplot depicts the median (center line), interquartile range (box limits, representing the first and third quartiles), and whiskers (extending to 1.5 × the interquartile range). **(C, D)** Similar plots to panels A and B, but for human-to-total (H:Total) ratios (normalized by total metagenomic reads from a sample). **(E, F)** Similar plots to panels A and B, but for bacterial-to-total (B:Total) ratios (raw p-values for the boxplots multiplied by six, which was the number of pairwise comparisons made, prior to applying the alpha < 0.05 threshold). For all panels: Bonferoni-corrected ***P < 0.0001, **P< 0.0016 (two-sided Welch’s t-test). The data were obtained from Poyet et al. (2019).

### Tracking response to antibiotic treatment with B:H ratios

We pulled down metagenomic data from two studies by Palleja et al. (2018) and Chng et al. (2020) that treated humans and mice with antibiotics, respectively, sampling before, during, and after treatment ^4,18^. We plotted the log-transformed B:H ratios from human stool metagenomic data sampled from 12 individuals across five time points: Day 0 (baseline), Day 4 (during antibiotic intervention), Day 8, Day 42, and Day 180 (post-antibiotic recovery; **Fig. 5A**). We observed a significant decline in B:H ratios from Day 0 to Day 4 (two-sided Welch’s T-test, p < 0.001), indicating significant bacterial biomass depletion due to antibiotics, followed by a rapid recovery to baseline levels by day 8, which persisted throughout the time series (**Fig. 5A**). We saw a similar pattern for log-transformed B:H ratios sampled from 27 mice across nine time points, with a steep drop in bacterial biomass during antibiotic treatment (Days 3, 6, and 7; two sided Welch’s T-test, p < 0.001), with ratios returning to baseline levels by Day 10 (**Fig. 5B**). Both analyses demonstrated consistent patterns of rapid microbiome depletion during antibiotic exposure, with an average 45-fold drop in B:H ratios in the human cohort and an average 403-fold drop in the B:H ratios in the mouse data, followed by recovery within several days. In the human cohort, B:H ratios differed cross-sectionally by less than 8-9 at baseline. Overall, we find that cross-sectional variation in B:H ratios in healthy human stool is less than 9-fold, while antibiotic-induced drops in B:H ratios are on the order of 45-fold.

**Figure 5.**
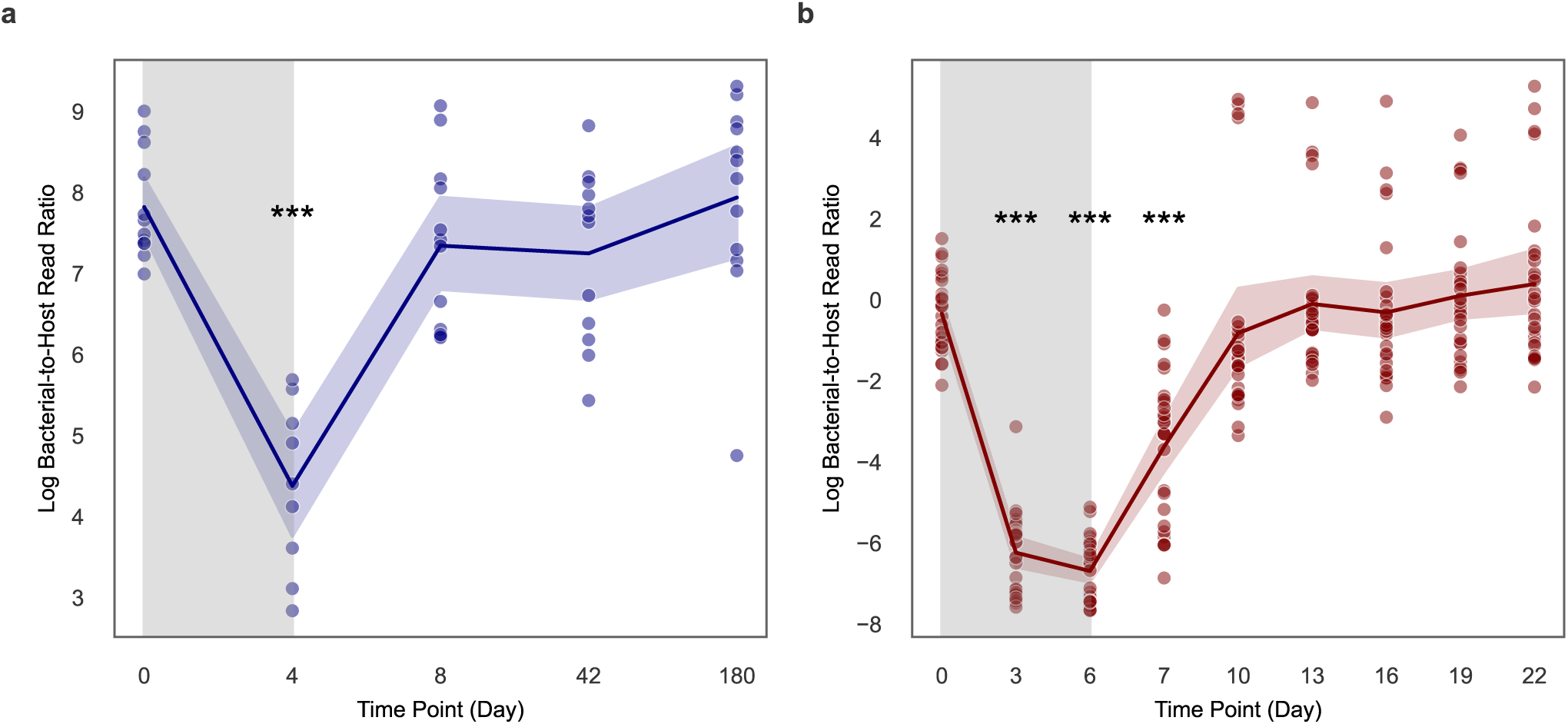
Temporal dynamics of bacteria-to-host ratios before, during, and after antibiotic treatment in humans and in mice. (**A**) Line plot showing the log-transformed B:H read ratios from metagenomic sequencing of human stools sampled from 12 individuals (n = 58) across five time points (days): baseline (day 0), during (day 4), and after (days 8, 42, and 180) antibiotic treatment. An initial sharp decline was observed, followed by a rapid recovery post-antibiotics. The data were obtained from Palleja et al. (2018). (**B**) Line plot showing the log-transformed B:H read ratios from metagenomic sequencing of mouse stools sampled from 27 mice (n = 243) over nine time points (days): baseline (day 0), during (days 3-6), and after (days 7, 10, 13, 16, 19, and 22) antibiotic exposure, showing a similar pattern of depletion and recovery. These data were derived from Chng et al. (2020). Blue and red shading around lines represent 95% confidence intervals, and the gray shaded regions indicate the antibiotic treatment windows.

## Discussion

In this study, we asked whether normalization by host reads alone was sufficient to estimate absolute bacterial biomass directly from stool metagenomic data, without the need for synthetic spike-ins or or additional experimental measurements. We compared and contrasted B:H ratios to other more established biomass estimation methods and we validated B:H ratios using longitudinal data from humans and mice treated with antibiotics.

We found that stool moisture content was inversely associated with cytometric cell counts per gram of fresh stool (i.e., often termed ‘microbial load’) ^30^. B:H ratios showed a weak positive association with microbial load and no association with Bristol stool scores (a proxy for water content), indicating that B:H ratios may be more direct measures of bacterial biomass (i.e., independent of stool consistency or moisture content; **Fig. 2**). Our finding is consistent with prior studies ^18,21,32^, which identified stool moisture content and bowel movement frequency, respectively, as major confounding factors in microbiome analyses. As we outline above, stool consistency is not necessarily related to total bacterial biomass in the gut (e.g., intuitively, a vegan with loose stool could have much higher total bacterial biomass in their gut than a carnivore with constipation, but this might not be apparent when looking at microbial load) ^30,33,34^, and it is important to have biomass estimates that are independent of moisture content and transit time.

We saw strong agreement between qPCR-based estimates of absolute 16S copy number and B:H read ratios (**Fig. 3A**). In the original study, the authors found that plant-derived reads in the metagenomic data, likely coming from the diet, were inversely related to qPCR biomass estimates. We found that normalizing by both plant-derived and host-derived reads provided roughly equivalent estimates of bacterial biomass (**Fig. 3B**). Together, these findings reinforce the utility of B:H ratios, and perhaps bacterial-to-dietary read ratios, for generating accurate estimates of bacterial biomass in the mammalian gut. Unlike mice, however, humans consume a wide variety of diets, and there is evidence that dietary read frequencies in human stool can fluctuate over several orders of magnitude, depending on the types of foods consumed ^35^. As such, host DNA may be a more reliable normalization factor in human stool.

Perhaps the most widely accepted method for biomass normalization in metagenomic sequencing is the spiking-in of a controlled amount of cells or DNA from an organism that is not present in the system (e.g., a hyperthermophile spiked into a stool sample). We looked at a largest synthetic spike-in data set we could identify (N=385), which consisted of cow milk samples where *ZymoBIOMICSTM Spike-in Control I (High Microbial Load)* was added in to assess bacterial biomass levels. We found that the fraction of host DNA was tightly associated with the abundance of the spike-in (r^2^ = 0.784; **Fig. 4**). Synthetic spike-ins require additional experimental design considerations and they reduce the number of sequencing reads from target organisms. Leveraging natural spike-ins, like host-derived sequences, appears to be sufficient for absolute biomass estimation. Taken together with the fecal samples above, this result suggests that host-associated substrates, like stool, milk, vaginal fluid, or saliva contain a relatively stable amount of host DNA that can be leveraged for bacterial absolute biomass estimation.

As a final assessment of our approach, we analyzed time series data from healthy human and mouse cohorts that received broad-spectrum antibiotic treatments (**Fig. 5A-B**). B:H ratios showed a 45-fold and a 403-fold drop following antibiotic treatment in humans and mice, respectively, followed by a recovery back to the baseline B:H level (**Fig. 5A-B**). These antibiotic-induced shifts in estimated biomass are much larger than the 2.4 fold difference observed between average B:H ratios across the Poyet et al. (2019) metagenomic time series and the 8-9 fold differences observed in the human antibiotic cohort at the baseline time point, indicating that cross-sectional variation in healthy individuals is substantially smaller than major, clinically-relevant disruptions to gut bacterial biomass (**Fig. 4A-B**). Major declines in gut bacterial biomass have been associated with inflammatory bowel disease, antibiotic treatment, chemotherapy treatment, and gastrointestinal cancers, while higher bacterial biomass and diversity have been associated with both health and constipation ^21,36^.

In conclusion, the B:H ratio represents a simple approach for estimating absolute bacterial biomass in stool, and possibly in other host-derived substrates, leveraging host read counts that are often disregarded in metagenomic sequencing studies. While the assumption that epithelial shedding is equivalent across humans is not strictly true, given that we observe, at most, a 9-fold variation across healthy individuals, the scale of cross-sectional variation is much smaller than antibiotic-induced biomass fluctuations in healthy individuals. Host-derived read normalization can be applied to existing metagenomic datasets without requiring additional measurements from conventional methods, which are typically resource-intensive, time-consuming, require specialized expertise, and suffer from several sources of noise and bias. The B:H ratio appears to be largely independent of stool consistency and water content, unlike most other stool bacterial biomass measures that are normalized per unit wet-weight. However, validation data are needed, with paired measures of microbial load and fecal water content, to assess how B:H ratios and bacterial biomass (per unit dry weight) vary across health and disease states. Absolute bacterial biomass is a key metric that often gets left out of gut microbiome studies, and empowering researchers to include this measure more broadly in their metagenomic analyses should serve to improve our understanding of host-microbiota interactions.

## Methods

### Data Sources and Processing

#### 16S Sequencing data from a study cohort of of 40 volunteers, 20 from a longitudinal cohort and 29 patients with Crohn’s disease and 66 healthy controls a disease cohort

Pre-processed data were taken directly from the supplementary information section of Vandeputte et al. (2017) ^18^. This dataset included bacterial biomass measurements, determined by a C6 Accuri flow cytometer (BD Biosciences) after mechanical homogenization, as well as stool moisture content that was measured in duplicate as the percentage of mass loss after freeze-drying 0.2 g of frozen, homogenized fecal material stored at −80°C.

#### Metagenomic data from the European Metacardis Cohort

Pre-processed data were obtained directly from supplementary tables (Tables 1-18) of Fromentin et al. (2022) ^25^. The Metacardis cohort included 869 healthy controls (HCs) and individuals across varying stages of dysmetabolism and ischemic heart disease (IHD) severity, aged 18–75 years, recruited from Denmark, France, and Germany between 2013 and 2015. Bacterial biomass in stool samples was quantified using a C6 Accuri flow cytometer and expressed as cell counts per gram of fecal material (i.e., microbial load index). Fecal DNA was extracted and sequenced, yielding an average of 23.3 million single-end short reads (± 4.0 million, s.d.) with a mean read length of 150 bases. Bacterial biomass, as well as bacterial, host, and total read counts were taken directly from the supplemental tables.

#### Gut Puzzle Manifest Metagenomic dataset

Fecal samples from 39 Gut Puzzle participants with Bristol Stool Score metadata were collected and processed using our lab’s custom Nextflow pipeline (https://github.com/Gibbons-Lab/pipelines/tree/master/metagenomics). Samples were collected in 1,200□ml 2-piece specimen collectors (Medline) in the Public Health Science Division of the Fred Hutchinson Cancer Center (IRB protocol number 10961) and transferred into a large vinyl anaerobic chamber (Coy; 37□°C, 5% hydrogen, 20% carbon dioxide, balanced with nitrogen) at the Institute for Systems Biology within 30□min of sample receipt. Fecal aliquots were sent to Diversigen, Inc., for DNA extraction, library preparation, and shotgun metagenomic sequencing. Briefly, libraries were prepared with the Nextera XT Library Prep kit (Illumina) and sequenced with a paired-end 2□×□150□bp protocol on a NovaSeq 6000 (Illumina) yielding at least 70□M reads per sample. Initial quality control was performed using fastp^37^, where reads were trimmed to remove low-quality bases, with a minimum quality threshold set at 20, a minimum read length of 50 bp, and a maximum read length of 150 bp to ensure the retention of high-quality data. Taxonomic relative abundances were estimated using Kraken2 ^38^ and Bracken ^39^, with a custom Kraken2 database (kraken2_db_uhgg_v2.0.1 database) constructed using data from Almeida et. al. (2020) ^40^, including the human genome. For this analysis, we used a confidence threshold of 0.3 for genus and species-level identification across multiple taxonomic ranks.

#### Mouse antibiotic treatment data set

Raw metagenomic data (FASTQ files) from Chng et al. (2020) ^4^ were downloaded from Sequence Read Archive (SRA) under the accession number SRP142225. The data were reprocessed using the same Nextflow-based pipeline described above, with a modified Kraken2 database specific to mouse metagenomes, where the human reference genome was replaced with the Genome Reference Consortium Mouse Build 39 (GRCm39) ^41^. Taxonomic abundances were estimated using Bracken ^39^, applying an abundance cutoff of 10 reads before reassignment. Stools were sampled as a cage unit (two mice per cage) over multiple time points: before antibiotic treatment (day 0), mid-point of antibiotic treatment (day 3), end-point of antibiotic treatment (day 6), 1-day post-gavage (day 7), 4-day post-gavage (day 10), 7-day post-gavage (day 13), 10-day post-gavage (day 16), 13-day post-gavage (day 19) and 16-day post-gavage (day 22). Total bacterial DNA was extracted from fecal samples using the PowerSoil DNA isolation kit (MoBio Laboratories) according to the manufacturer’s instructions.

Absolute 16S rRNA genes were quantified with qPCR using a pair of universal 16S primers. DNA from six treatment groups was amplified on days 0, 3, 10, and 13. Each reaction was prepared in triplicate on a 384-well plate, containing 5 µl PowerUp SYBR Green Master Mix, 0.5□µl of 5□µM primers and 1□µl of 10× diluted DNA, with a total volume of 10 µl. The ViiA 7 Real-Time PCR System (Thermo Fisher Scientific) was used for qPCR with the following amplification specifications: 1 cycle of 95□°C for 2□min, 40 cycles of 95□°C for 15□s, 60□°C for 15□s, and 72□°C for 1□min. A standard curve, created from serial dilutions of synthesized DNA, was used to convert Ct values to copy numbers, and day 0 copy numbers were used to normalize bacterial abundances across samples. These results were directly sourced from the supplement.

Diet-normalized bacterial biomass was estimated by normalizing all reads classified to bacterial taxa with plant-derived reads on the assumption that the amount of diet-derived plant DNA would be conserved across mouse fecal samples. These data were sourced directly from the supplement.

#### Metagenomic data of bulk milk samples

Processed metagenomic data from cow milk samples, including total bacterial reads, total host reads, and total spike-in reads were obtained from the supplementary materials section of Wallace et al. (2023) ^16^. Bulk milk samples were collected from 276 commercial dairy cows in New Zealand. For these bulk samples, a 15 mL subsample was sent weekly to the Herd Testing facility for host cell counting. All samples were stored at -20°C prior to DNA extraction and spiked with 17 µL of a 1:100 diluted spike-in control (ZymoBIOMICS™ Spike-in Control I, High Microbial Load). Following extraction, short-read shotgun sequencing libraries (150 bp paired-end) were prepared using the Illumina DNA Flex library prep kit, and sequencing was performed on an Illumina NovaSeq system with S1 and S4 flowcells, aiming for 15 million reads per sample. Quality control checks were conducted using the FastQC program, and samples with fewer than 100,000 reads were excluded from further analysis. Reads were classified using Kraken2, against a database comprising microbiome, human, and bovine sequences downloaded from NCBI’s RefSeq database, allowing identification of bacterial, host (cow genome), and spike-in reads.

#### Metagenomic data of a cohort of 12 healthy humans given a four-day antibiotic intervention

Pre-processed metagenomic data, including total bacterial read counts, host read counts, and overall read counts, were obtained from the supplementary information of Palleja et al. (2018) ^6^. Stool samples were collected from 12 healthy Caucasian men who were 18 to 40 years of age. In addition to a screening visit, the study design encompassed five study visits (D0, D4, D8, D42 and D180) and a four-day broad-spectrum antibiotic intervention consisting of once-daily administration of 500□mg meropenem, 500□mg vancomycin and 40□mg gentamicin dissolved in apple juice and ingested orally. Microbial DNA was extracted from 200□mg frozen stool and sequenced. An average of 79.4 ± 18.0 million raw metagenomic reads per sample were generated, corresponding to 7.94 ± 1.8 Gb of data. The average sequencing depths for samples collected at time points D0, D4, D8, D42, and D180 were 76.5 ± 11.1, 78.1 ± 13.2, 75.6 ± 19.6, 81.2 ± 11.4, and 85.4 ± 26.6 million reads, respectively, indicating no significant reduction in read depths immediately following the intervention. To ensure data quality, reads were subjected to adaptor removal and trimmed based on a quality score threshold of 20 and a minimum read length of 30 base pairs. This process resulted in an average of 6.8 million reads being discarded due to adaptor contamination, while 0.94 million reads were removed for not meeting the trimming criteria. Human DNA contamination was eliminated by aligning reads against the human genome (version hg19). Reads excluded during this step were, approximately 0.24 million reads per sample were used to calculate the host read counts for subsequent analysis. After these quality control measures, the final datasets contained high-quality non-human reads of 69.3 ± 8.7 million for D0, 66.5 ± 13.1 million for D4, 68.2 ± 15.8 million for D8, 72.0 ± 12.1 million for D42, and 79.8 ± 22.6 million for D180.

#### Longitudinal BIO-ML metagenomic data of a cohort of 4 healthy individuals

The raw metagenomic data (FASTQ files), including total bacterial read counts, host read counts, and overall read counts, belonging to Poyet et al. (2019) ^31^, was downloaded from Sequence Read Archive (SRA) under the accession number PRJNA544527 ^31^ and reprocessed using the same Nextflow-based pipeline as described above. A total of 1,207 stool samples were collected from 90 participants between July 2014 and May 2016 and sourced from the non-profit stool bank OpenBiome. Donors, aged 19 to 45 years (mean age of 28), had body mass indexes from 17.5 to 29.8 (mean of 23.4) and were screened by OpenBiome to ensure they were healthy and pathogen-free. Samples were deidentified, diluted 1:10 in a solution of 12.5% glycerol and 0.9% NaCl, homogenized, and filtered through a 330-μm filter. DNA was extracted using the MoBio PowerSoil 96 kit (Qiagen Cat No. 12955-4) with minor modifications. After thawing on ice, 625 μL to 1 mL of homogenized stool was added to the PowerSoil plate (12955-4-BP) and centrifuged at 4,000g for 10 minutes. Following removal of the supernatant, 750 μL of bead solution and 60 μL of C1 solution were added. Samples were bead-beaten at 20 Hz for 10 minutes, rotated 180 degrees, and beaten for an additional 10 minutes. They were then centrifuged at 4,500g for 6 minutes, and 850 μL of the supernatant was transferred to a clean collection plate. The remaining steps followed the manufacturer’s protocol. Metagenomic DNA was quantified using the Quant-iT PicoGreen dsDNA Assay (Life Technologies) and normalized to 50 pg/μL. Illumina sequencing libraries were generated from 100–250 pg of DNA with the Nextera XT DNA Library Preparation kit (Illumina), following the manufacturer’s protocol with scaled reaction volumes. Libraries were pooled by combining 200 nl from each of 96 samples. Insert sizes and concentrations of the pooled libraries were verified with an Agilent Bioanalyzer DNA 1000 kit (Agilent Technologies). Sequencing was performed on a HiSeq system (2 × 101 bp), targeting ∼10 million paired-end reads. Shotgun metagenomic sequencing data underwent quality trimming to remove low-quality bases and human-aligned reads (hg19), followed by duplicate sequence removal using fastuniq. This process yielded approximately 9.8 × 10□high-quality reads per sample. The filtered reads were assembled with metaSPAdes, and protein-coding genes were identified using Prodigal. To reduce redundancy, genes were clustered with CD-HIT to generate a nonredundant gene set, which was subsequently annotated with COG terms using rps-blast. Finally, Bowtie2 was employed to align the reads to the COG-annotated gene set, and the relative abundances of COG families were determined based on gene coverage.

### Statistical Analyses

Due to the right-skewed nature of the B:H ratios, bacteria-to-plant ratios, bacteria-to-spike-in ratios, and qPCR copy numbers, we applied a natural log transformation so that distributions behaved more normally. Pairwise comparisons of the log-transformed B:H, B:Total, and H:Total ratios among the four donors in the dense time-series analyses were conducted using Welch’s t-tests, assuming unequal variance. A Bonferroni correction was applied to adjust for multiple testing, with the significance threshold set to □= 0.05/N, where N is the number of pairwise comparisons. Corrected p-values were evaluated against this adjusted threshold. Linear regressions were used to assess the relationships between the log-transformed B:H ratios and other bacterial biomass metrics, with the B:H ratios serving as the independent variable. Ordinal logistic regression model was used to examine the relationship between stool consistency (Bristol score) and the B:H ratios, with consistency as the dependent variable. Additionally, line plots were employed to visualize the distributions of log bacterial-to-host read ratios in humans and compare them with bacterial-to-host read ratios in mice. All statistical analyses and visualizations were conducted using Python 3.9.19 with the following libraries: pandas (1.5.3), numpy (1.26.4), statsmodels (0.14.0), matplotlib (3.8.4), seaborn (0.13.2), scipy (1.12.0), and scikit-learn (1.2.2). See code availability section for analysis notebooks.

## Data availability

Preprocessed 16S amplicon data from Vandepuette et al. (2017) are available in the supplementary information section (Tables 1-11). Preprocessed metagenomic data from Fromentin et al. (2022) are available in the supplementary information section (Tables 1-18). Preprocessed metagenomic data of milk samples by Wallace et al. (2023) are provided in the supplementary data tables of the supplementary material section. Raw metagenomic data from Chng et al. (2020) can be found in the Sequence Read Archive (SRA) under project ID SRP142225. Raw metagenomic data from Palleja et al. (2018) are accessible in the European Nucleotide Archive (ENA) under accession number ERP022986. Raw metagenomic data from Poyet et al. (2019) can be found at NCBI BioProject under accession number PRJNA544527. Gut Puzzle data will be uploaded to SRA prior to publication of this work.

## Code availability

Nextflow pipelines for processing metagenomic shotgun sequencing data, from raw reads to taxonomic abundance matrices, are available https://github.com/Gibbons-Lab/pipelines/ (metagenomics pipeline). Scripts used for analyzing the data and generating the figures in this study can be accessed at https://github.com/Gibbons-Lab/Metagenomic_Biomass_Quantification_2024

## Acknowledgements

Research reported in this publication was supported by the National Institute of Diabetes and Digestive and Kidney Diseases (NIDDK) of the National Institutes of Health (NIH) under award number R01DK133468 (to SMG). The faecal sample collection at Fred Hutchinson Cancer Center for the Gut Puzzle study was supported by P30 CA015704.

## Supplementary Information

**Figure S1.**
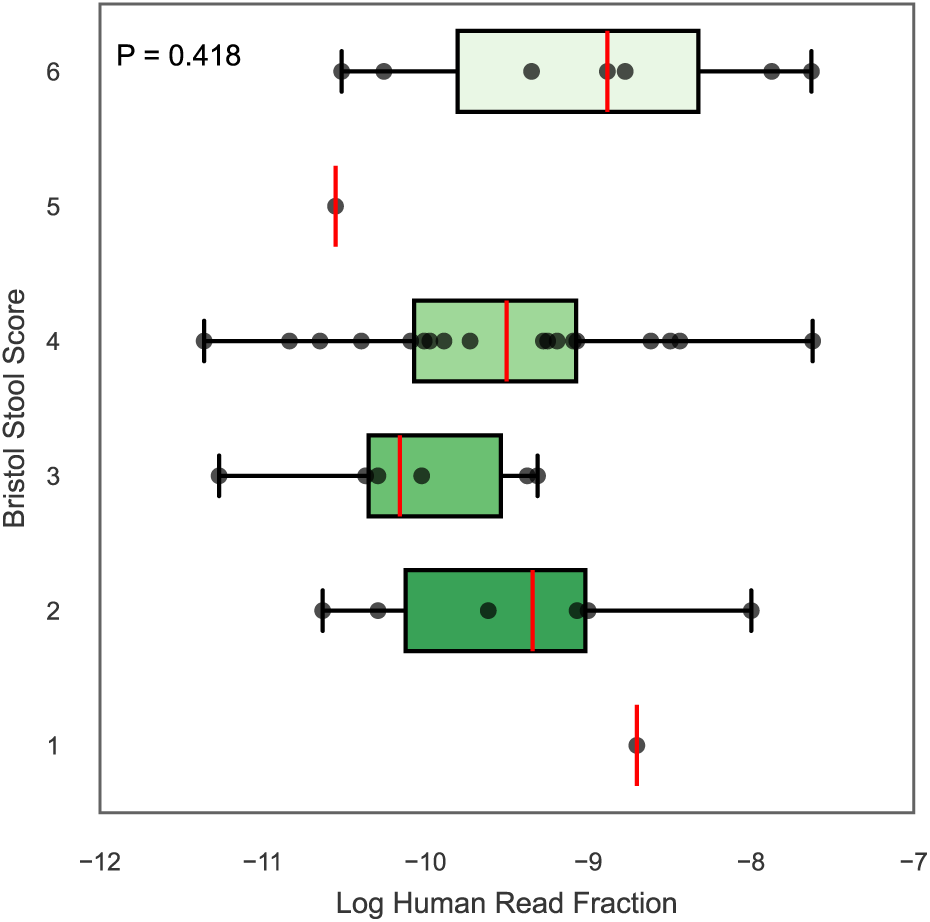
Distribution of log-transformed human read fractions across different Bristol Stool Scores. Boxplots showing human read fractions (relative to total metagenic reads) across Bristol stool score categories (n = 39). Each boxplot displays the center line (median), box limits (first and third quartiles), and whiskers (1.5 × interquartile range). Using ordinal logistic regression, we did not observe a significant association between human read fractions and Bristol scores (P=0.418).

## References

1. Human Microbiome Project Consortium. Structure, function and diversity of the healthy human microbiome. Nature 486, 207–214 (2012).

2. Wang, W.-L., Xu, S.-Y., Ren, Z.-G., Tao, L., Jiang, J.-W. & Zheng, S.-S. Application of metagenomics in the human gut microbiome. World J. Gastroenterol. 21, 803–814 (2015).

3. Rose, C., Parker, A., Jefferson, B. & Cartmell, E. The Characterization of Feces and Urine: A Review of the Literature to Inform Advanced Treatment Technology. Crit. Rev. Environ. Sci. Technol. 45, 1827–1879 (2015).

4. Chng, K. R., Ghosh, T. S., Tan, Y. H., Nandi, T., Lee, I. R., Ng, A. H. Q., Li, C., Ravikrishnan, A., Lim, K. M., Lye, D., Barkham, T., Raman, K., Chen, S. L., Chai, L., Young, B., Gan, Y.-H. & Nagarajan, N. Metagenome-wide association analysis identifies microbial determinants of post-antibiotic ecological recovery in the gut. Nature Ecology & Evolution 4, 1256–1267 (2020).

5. Diener, C., Dai, C. L., Wilmanski, T., Baloni, P., Smith, B., Rappaport, N., Hood, L., Magis, A. T. & Gibbons, S. M. Genome-microbiome interplay provides insight into the determinants of the human blood metabolome. Nat Metab 4, 1560–1572 (2022).

6. Palleja, A., Mikkelsen, K. H., Forslund, S. K., Kashani, A., Allin, K. H., Nielsen, T., Hansen, T. H., Liang, S., Feng, Q., Zhang, C., Pyl, P. T., Coelho, L. P., Yang, H., Wang, J., Typas, A., Nielsen, M. F., Nielsen, H. B., Bork, P., Wang, J., Vilsbøll, T., Hansen, T., Knop, F. K., Arumugam, M. & Pedersen, O. Recovery of gut microbiota of healthy adults following antibiotic exposure. Nat Microbiol 3, 1255–1265 (2018).

7. Kurilshikov, A., Medina-Gomez, C., Bacigalupe, R., Radjabzadeh, D., Wang, J., Demirkan, A., Le Roy, C. I., Raygoza Garay, J. A., Finnicum, C. T., Liu, X., Zhernakova, D. V., Bonder, M. J., Hansen, T. H., Frost, F., Rühlemann, M. C., Turpin, W., Moon, J.-Y., Kim, H.-N., Lüll, K., Barkan, E., Shah, S. A., Fornage, M., Szopinska-Tokov, J., Wallen, Z. D., Borisevich, D., Agreus, L., Andreasson, A., Bang, C., Bedrani, L., Bell, J. T., Bisgaard, H., Boehnke, M., Boomsma, D. I., Burk, R. D., Claringbould, A., Croitoru, K., Davies, G. E., van Duijn, C. M., Duijts, L., Falony, G., Fu, J., van der Graaf, A., Hansen, T., Homuth, G., Hughes, D. A., Ijzerman, R. G., Jackson, M. A., Jaddoe, V. W. V., Joossens, M., Jørgensen, T., Keszthelyi, D., Knight, R., Laakso, M., Laudes, M., Launer, L. J., Lieb, W., Lusis, A. J., Masclee, A. A. M., Moll, H. A., Mujagic, Z., Qibin, Q., Rothschild, D., Shin, H., Sørensen, S. J., Steves, C. J., Thorsen, J., Timpson, N. J., Tito, R. Y., Vieira-Silva, S., Völker, U., Völzke, H., Võsa, U., Wade, K. H., Walter, S., Watanabe, K., Weiss, S., Weiss, F. U., Weissbrod, O., Westra, H.-J., Willemsen, G., Payami, H., Jonkers, D. M. A. E., Arias Vasquez, A., de Geus, E. J. C., Meyer, K. A., Stokholm, J., Segal, E., Org, E., Wijmenga, C., Kim, H.-L., Kaplan, R. C., Spector, T. D., Uitterlinden, A. G., Rivadeneira, F., Franke, A., Lerch, M. M., Franke, L., Sanna, S., D’Amato, M., Pedersen, O., Paterson, A. D., Kraaij, R., Raes, J. & Zhernakova, A. Large-scale association analyses identify host factors influencing human gut microbiome composition. Nat. Genet. 53, 156–165 (2021).

8. Rothschild, D., Weissbrod, O., Barkan, E., Kurilshikov, A., Korem, T., Zeevi, D., Costea, P. I., Godneva, A., Kalka, I. N., Bar, N., Shilo, S., Lador, D., Vila, A. V., Zmora, N., Pevsner-Fischer, M., Israeli, D., Kosower, N., Malka, G., Wolf, B. C., Avnit-Sagi, T., Lotan-Pompan, M., Weinberger, A., Halpern, Z., Carmi, S., Fu, J., Wijmenga, C., Zhernakova, A., Elinav, E. & Segal, E. Environment dominates over host genetics in shaping human gut microbiota. Nature 555, 210–215 (2018).

9. Manor, O., Dai, C. L., Kornilov, S. A., Smith, B., Price, N. D., Lovejoy, J. C., Gibbons, S. M. & Magis, A. T. Health and disease markers correlate with gut microbiome composition across thousands of people. Nat. Commun. 11, 5206 (2020).

10. Gloor, G. B., Macklaim, J. M., Pawlowsky-Glahn, V. & Egozcue, J. J. Microbiome Datasets Are Compositional: And This Is Not Optional. Front. Microbiol. 8, 2224 (2017).

11. Barlow, J. T., Bogatyrev, S. R. & Ismagilov, R. F. A quantitative sequencing framework for absolute abundance measurements of mucosal and lumenal microbial communities. Nat. Commun. 11, 2590 (2020).

12. Tito, R. Y., Verbandt, S., Aguirre Vazquez, M., Lahti, L., Verspecht, C., Lloréns-Rico, V., Vieira-Silva, S., Arts, J., Falony, G., Dekker, E., Reumers, J., Tejpar, S. & Raes, J. Microbiome confounders and quantitative profiling challenge predicted microbial targets in colorectal cancer development. Nat. Med. 30, 1339–1348 (2024).

13. Vandeputte, D., De Commer, L., Tito, R. Y., Kathagen, G., Sabino, J., Vermeire, S., Faust, K. & Raes, J. Temporal variability in quantitative human gut microbiome profiles and implications for clinical research. Nat. Commun. 12, 6740 (2021).

14. Stämmler, F., Gläsner, J., Hiergeist, A., Holler, E., Weber, D., Oefner, P. J., Gessner, A. & Spang, R. Adjusting microbiome profiles for differences in microbial load by spike-in bacteria. Microbiome 4, 28 (2016).

15. Morgan, X. C. & Huttenhower, C. Meta’omic analytic techniques for studying the intestinal microbiome. Gastroenterology 146, 1437–1448.e1 (2014).

16. Wallace, A., Ling, H., Gatenby, S., Pruden, S., Neeley, C., Harland, C. & Couldrey, C. Absolute Quantification of Microbiota in Shotgun Sequencing Using Host Cells or Spike-Ins. bioRxiv 2023.08.23.554046 (2023). doi:10.1101/2023.08.23.554046

17. Galazzo, G., van Best, N., Benedikter, B. J., Janssen, K., Bervoets, L., Driessen, C., Oomen, M., Lucchesi, M., van Eijck, P. H., Becker, H. E. F., Hornef, M. W., Savelkoul, P. H., Stassen, F. R. M., Wolffs, P. F. & Penders, J. How to Count Our Microbes? The Effect of Different Quantitative Microbiome Profiling Approaches. Front. Cell. Infect. Microbiol. 10, 551454 (2020).

18. Vandeputte, D., Kathagen, G., D’hoe, K., Vieira-Silva, S., Valles-Colomer, M., Sabino, J., Wang, J., Tito, R. Y., De Commer, L., Darzi, Y., Vermeire, S., Falony, G. & Raes, J. Quantitative microbiome profiling links gut community variation to microbial load. Nature 551, 507–511 (2017).

19. Smith, C. J. & Osborn, A. M. Advantages and limitations of quantitative PCR (Q-PCR)-based approaches in microbial ecology. FEMS Microbiol. Ecol. 67, 6–20 (2009).

20. Tourlousse, D. M., Yoshiike, S., Ohashi, A., Matsukura, S., Noda, N. & Sekiguchi, Y. Synthetic spike-in standards for high-throughput 16S rRNA gene amplicon sequencing. Nucleic Acids Res. 45, e23 (2017).

21. Nishijima, S., Stankevic, E., Aasmets, O., Schmidt, T. S. B., Nagata, N., Keller, M. I., Ferretti, P., Juel, H. B., Fullam, A., Robbani, S. M., Schudoma, C., Hansen, J. K., Holm, L. A., Israelsen, M., Schierwagen, R., Torp, N., Telzerow, A., Hercog, R., Kandels, S., Hazenbrink, D. H. M., Arumugam, M., Bendtsen, F., Brøns, C., Fonvig, C. E., Holm, J.-C., Nielsen, T., Pedersen, J. S., Thiele, M. S., Trebicka, J., Org, E., Krag, A., Hansen, T., Kuhn, M., Bork, P. & GALAXY and MicrobLiver Consortia. Fecal microbial load is a major determinant of gut microbiome variation and a confounder for disease associations. Cell (2024). doi:10.1016/j.cell.2024.10.022

22. Morton, J. T., Marotz, C., Washburne, A., Silverman, J., Zaramela, L. S., Edlund, A., Zengler, K. & Knight, R. Establishing microbial composition measurement standards with reference frames. Nat Commun 10, 2719 (2019).

23. Vincent, C., Mehrotra, S., Loo, V. G., Dewar, K. & Manges, A. R. Excretion of Host DNA in Feces Is Associated with Risk of Clostridium difficile Infection. J Immunol Res 2015, 246203 (2015).

24. Shi, Y., Wang, G., Lau, H. C.-H. & Yu, J. Metagenomic Sequencing for Microbial DNA in Human Samples: Emerging Technological Advances. Int. J. Mol. Sci. 23, (2022).

25. Fromentin, S., Forslund, S. K., Chechi, K., Aron-Wisnewsky, J., Chakaroun, R., Nielsen, T., Tremaroli, V., Ji, B., Prifti, E., Myridakis, A., Chilloux, J., Andrikopoulos, P., Fan, Y., Olanipekun, M. T., Alves, R., Adiouch, S., Bar, N., Talmor-Barkan, Y., Belda, E., Caesar, R., Coelho, L. P., Falony, G., Fellahi, S., Galan, P., Galleron, N., Helft, G., Hoyles, L., Isnard, R., Le Chatelier, E., Julienne, H., Olsson, L., Pedersen, H. K., Pons, N., Quinquis, B., Rouault, C., Roume, H., Salem, J.-E., Schmidt, T. S. B., Vieira-Silva, S., Li, P., Zimmermann-Kogadeeva, M., Lewinter, C., Søndertoft, N. B., Hansen, T. H., Gauguier, D., Gøtze, J. P., Køber, L., Kornowski, R., Vestergaard, H., Hansen, T., Zucker, J.-D., Hercberg, S., Letunic, I., Bäckhed, F., Oppert, J.-M., Nielsen, J., Raes, J., Bork, P., Stumvoll, M., Segal, E., Clément, K., Dumas, M.-E., Ehrlich, S. D. & Pedersen, O. Microbiome and metabolome features of the cardiometabolic disease spectrum. Nat. Med. 28, 303–314 (2022).

26. Yu, L., Jia, R., Liu, S., Li, S., Zhong, S., Liu, G., Zeng, R. J., Rensing, C. & Zhou, S. Ferrihydrite-mediated methanotrophic nitrogen fixation in paddy soil under hypoxia. ISME Commun 4, ycae030 (2024).

27. Maxfield, P. J., Hornibrook, E. R. C. & Evershed, R. P. Estimating high-affinity methanotrophic bacterial biomass, growth, and turnover in soil by phospholipid fatty acid 13C labeling. Appl. Environ. Microbiol. 72, 3901–3907 (2006).

28. Aichbichler, B. W., Wenzl, H. H., Santa Ana, C. A., Porter, J. L., Schiller, L. R. & Fordtran, J. S. A comparison of stool characteristics from normal and constipated people. Dig. Dis. Sci. 43, 2353–2362 (1998).

29. Blake, M. R., Raker, J. M. & Whelan, K. Validity and reliability of the Bristol Stool Form Scale in healthy adults and patients with diarrhoea-predominant irritable bowel syndrome. Aliment. Pharmacol. Ther. 44, 693–703 (2016).

30. Vandeputte, D., Falony, G., Vieira-Silva, S., Tito, R. Y., Joossens, M. & Raes, J. Stool consistency is strongly associated with gut microbiota richness and composition, enterotypes and bacterial growth rates. Gut 65, 57–62 (2016).

31. Poyet, M., Groussin, M., Gibbons, S. M., Avila-Pacheco, J., Jiang, X., Kearney, S. M., Perrotta, A. R., Berdy, B., Zhao, S., Lieberman, T. D., Swanson, P. K., Smith, M., Roesemann, S., Alexander, J. E., Rich, S. A., Livny, J., Vlamakis, H., Clish, C., Bullock, K., Deik, A., Scott, J., Pierce, K. A., Xavier, R. J. & Alm, E. J. A library of human gut bacterial isolates paired with longitudinal multiomics data enables mechanistic microbiome research. Nat Med 25, 1442–1452 (2019).

32. Johnson-Martínez, J. P., Diener, C., Levine, A. E., Wilmanski, T., Suskind, D. L., Ralevski, A., Hadlock, J., Magis, A. T., Hood, L., Rappaport, N. & Gibbons, S. M. Aberrant bowel movement frequencies coincide with increased microbe-derived blood metabolites associated with reduced organ function. Cell Rep Med 5, 101646 (2024).

33. Gilbert, J. A. & Alverdy, J. Stool consistency as a major confounding factor affecting microbiota composition: an ignored variable? Gut 65, 1–2 (2016).

34. Park, G., Kim, S., Lee, W., Kim, G. & Shin, H. Deciphering the Impact of Defecation Frequency on Gut Microbiome Composition and Diversity. Int. J. Mol. Sci. 25, 4657 (2024).

35. Diener, C. & Gibbons, S. M. Metagenomic estimation of dietary intake from human stool. bioRxiv (2024). doi:10.1101/2024.02.02.578701

36. DeGruttola, A. K., Low, D., Mizoguchi, A. & Mizoguchi, E. Current Understanding of Dysbiosis in Disease in Human and Animal Models. Inflamm. Bowel Dis. 22, 1137–1150 (2016).

37. Chen, S., Zhou, Y., Chen, Y. & Gu, J. fastp: an ultra-fast all-in-one FASTQ preprocessor. Bioinformatics 34, i884–i890 (2018).

38. Wood, D. E., Lu, J. & Langmead, B. Improved metagenomic analysis with Kraken 2. Genome Biology 20, 1–13 (2019).

39. Lu, J., Breitwieser, F. P., Thielen, P. & Salzberg, S. L. Bracken: estimating species abundance in metagenomics data. PeerJ Comput. Sci. 3, e104 (2017).

40. Almeida, A., Nayfach, S., Boland, M., Strozzi, F., Beracochea, M., Shi, Z. J., Pollard, K. S., Sakharova, E., Parks, D. H., Hugenholtz, P., Segata, N., Kyrpides, N. C. & Finn, R. D. A unified catalog of 204,938 reference genomes from the human gut microbiome. Nature Biotechnology 39, 105–114 (2020).

41. Kane, A. E., Chellappa, K., Schultz, M. B., Arnold, M., Li, J., Amorim, J., Diener, C., Zhu, D., Mitchell, S. J., Griffin, P., Tian, X., Petty, C., Conway, R., Walsh, K., Shelerud, L., Duesing, C., Mueller, A., Li, K., McNamara, M., Shima, R. T., Mitchell, J., Bonkowski, M. S., de Cabo, R., Gibbons, S. M., Wu, L. E., Ikeno, Y., Baur, J. A., Rajman, L. & Sinclair, D. A. Long-term NMN treatment increases lifespan and healthspan in mice in a sex dependent manner. bioRxivorg (2024). doi:10.1101/2024.06.21.599604

